# Development of first linkage map for *Silphium integrifolium* (*Asteraceae*) enables identification of sporophytic self-incompatibility locus

**DOI:** 10.1101/2021.01.29.428840

**Authors:** John H. Price, Andrew R. Raduski, Yaniv Brandvain, David L. Van Tassel, Kevin P. Smith

## Abstract

- *Silphium integrifolium* (*Asteraceae*) has been identified as a candidate for domestication as a perennial oilseed crop and has a sporophytic self-incompatibility system—the genetic basis of which is not well understood in the *Asteraceae*. To address this gap, we sought to map the genomic location of the self-recognition locus (S-locus) in this species.
- We used a biparental population and genotyping-by-sequencing to create the first genetic linkage map for this species. Then we developed a novel crossing scheme and set of analysis methods in order to infer S-locus genotypes for a subset of these individuals, allowing us to map the trait. Finally, we identified potential gene candidates using synteny analysis with the annual sunflower (*Helianthus annuus*) genome.
- Our linkage map contains 198 SNP markers and resolved into the correct number of linkage groups. We were able to successfully map the S-locus and identify several potential gene candidates in the sunflower syntenic region.
- Our method is effective and efficient, allowed us to map the *S. integrifolium* S-locus using fewer resources than previous studies, and could be readily be applied to other species. Our best gene candidate appears to be worthy of future work in *S. integrifolium* and other *Asteraceae* species.

## INTRODUCTION

*Silphium integrifolium* (Michx.) (wholeleaf rosinweed or silflower) is a member of the *Asteraceae* family native to prairies throughout the central United States. In the early 2000s, *S. integrifolium* was selected to be a candidate for domestication as a perennial oilseed crop by the Land Institute in Salina, Kansas (Van Tassel *et al*., 2017), attracting attention for its tolerance to drought, upright growth habit, and large seeds (DeHaan *et al*., 2016). Subsequently, *S. integrifolium* has been found to have a seed oil composition similar to landrace sunflower (Reinert et al., 2019) and good winter survival and persistence in a range of climates (J.H. Price & D.L. Van Tassel, pers. obs.). In addition, the yield potential of *S. integrifolium* populations that have undergone relatively little selection is approximately 60% the yield of advanced sunflower hybrids (Kandel *et al*., 2019; Schiffner *et al*., 2020), indicating that significant improvement is likely with continued breeding efforts. These characteristics further encourage the domestication of this species as a new crop. In the past, domestication occurred over long periods of time, largely due to the largely unintentional nature of early selection (Rindos, 1984). With the advantage of contemporary knowledge of genetics, genomics, and breeding techniques, the amount of time necessary to domesticate a new crop could be drastically reduced. Therefore, the development of genomic resources is a crucial step in this process (Sedbrook *et al*., 2014). To this end, we have developed the first genetic linkage map for *S. integrifolium*.

Among the traits for which better genetic knowledge will accelerate the domestication of *S. integrifolium* is self-incompatibility. Although occasional *S. integrifolium* individuals have been observed to produce at least some seed when self-pollinated (Reinert *et al*., 2020), *S. integrifolium* is self-incompatible, and as a member of the *Asteraceae* family is assumed to have a sporophytic self-incompatibility (SSI) system (Hiscock, 2000). In sporophytic systems, self-recognition is typically controlled by a single multi-allelic locus, known as the “S-locus”, with rejection of self-pollen caused by stigma recognition of S-locus gene products found in or on the pollen. Because these products are produced in the anther, pollen acceptance or rejection is determined by the diploid genotype of the male parent, rather than the haploid genotype of a given pollen grain (Hiscock & Tabah, 2003). SSI alleles are also expected to display complex dominance patterns, and dominance relationships between alleles may differ from the anther to the stigma (Hiscock & Tabah, 2003).

The molecular mechanisms that underlie SSI are best described in the *Brassicaceae*, where the female S-phenotype is determined by a receptor kinase complex known as SRK. When pollen of the same S-phenotype lands on the stigma, this kinase binds a cysteine-rich protein (CRP) found in the pollen coat, known as SP_11_/SCR, initiating the pollen rejection response (Fujii & Takayama, 2018). These two genes are tightly linked and rarely recombine; thus, they combine to form the S-locus, which is more properly thought of as an S-haplotype (Edh *et al.*, 2009). SRK is unable to bind SP_11_/SCR proteins produced by different haplotypes, resulting in the acceptance of non-self pollen (Fujii & Takayama, 2018). Although the identity of the receptor protein varies, secreted CRPs are also considered a promising candidate for the male determinant of SSI in the *Convolvulaceae* (morning glory) family (Rahman *et al*., 2007), and serve as the female determinant of gametophytic self-incompatibility (GSI) in the genus *Papaver* (poppy) (Marshall *et al*., 2011),.

*Asteraceae* systems are less well understood. In *Senecio squalidis*, and subsequently in other *Asteraceae*, SRK-like sequences have been identified and cloned. However, results from *S. squalidis* and chicory (*Cichorium intybus*) indicate that they likely are not integral to S-genotype determination (Hiscock & Tabah, 2003; Gonthier *et al*., 2013), and that the molecular control of *Asteraceae* SSI is quite different from the *Brassicaceae* system (Allen *et al.* 2011). Efforts to map the S-locus in chicory provided a 1.8 cM QTL region but have not yet determined a molecular basis for self-incompatibility (Gonthier *et al.*, 2013). Although efforts have been undertaken in other species, chicory represents perhaps the only example where a true *Asteraceae* S-locus has been definitively mapped, as opposed to other loci contributing to breakdowns in self-incompatibility (Gandhi *et al*., 2005; Koseva *et al*., 2017).

Mapping the S-locus is important for breeding efforts and for our understanding of the genetics and evolution of this critical locus. For example, the “collaborative nonself recognition” system identified in the *Solanaceae* (Kubo *et al.* 2011) has revealed that the mechanism underlying self-incompatibility can change how new S-alleles originate (Bod’ová *et al*., 2018, Harkness *et al*., 2021) and migrate (Harkness & Brandvain, 2020) across populations. Identifying the basis (or bases) of SSI in the *Asteraceae* would help us better understand and predict features of its evolution, including its maintenance in populations with very few (two to six) S-alleles (Brennan *et al*. 2006).

To map the S-locus, researchers must determine the S-locus genotype for individuals in a population large enough to conduct linkage mapping. In other species, S-genotype determination required mating large numbers of full-sibling individuals to one or a few “tester” genotypes with a known S-genotype (Camargo *et al*., 1997; Tomita *et al*., 2004). Testers may be a parent of the population, or may be obtained by mating siblings in a diallel design, grouping individuals based on their compatibility (Hiscock, 2000), and then selecting one or several of these individuals as a tester for their siblings (Gonthier *et al.*, 2013). However, this process has several limitations. For some species, pollen availability may limit the number of testcrosses that can be made, especially if clonal propagation is not used to multiply tester individuals. Additionally, because the tester is typically used as the male parent, differentiation between alleles may be difficult for SSI as dominance relationship between alleles may differ from anthers to stigma. Finally, this process often requires a large number of crosses, especially if the parents are not available. A diallel requires n(n-1) crosses, where **n** is the number of individuals to be analyzed, and multiple crosses are likely required for each individual being tested. Even selecting a tester from amongst a conservative number of siblings may require a prohibitively high number of crosses. Methods of S-allele determination that address these issues may make S-locus mapping feasible for a greater number of species.

In this study, we present a putative location for the *S. integrifolium* S-locus. To achieve this, we developed a novel framework for inferring the S-genotypes of individuals within a population large enough for mapping. Our method does not require tester individuals with known S-genotypes, requires only three to four crosses per individual, and identifies alleles with sex-specific dominance interactions. We used these inferred genotypes, in conjunction with the aforementioned linkage map, to identify a QTL likely containing the S-locus. We then identify regions in the sunflower (*Helianthus annuu*s) genome assembly syntenic with our putative S-locus and use this information to identify potential gene candidates for S-allele determination.

## MATERIALS AND METHODS

### Population development

A linkage map was constructed using an F_1_ population of 265 *Silphium integrifolium* (Michx.) individuals, derived from the crossing of two genotypes known as “965” and “1767”, selected at the Land Institute. These parents were chosen because they expressed agronomically favorable morphologies and differed for several phenological traits. Seventy-four seeds from this cross were grown at the Land Institute, and 191 were grown at the University of Minnesota. All 265 individuals were used for map construction, but only a subset of the 191 Minnesota individuals were used for S-allele determination. Following approximately two months of growth in a greenhouse, the Minnesota seedlings were transplanted to a field at the Minnesota Agricultural Experiment Station at the University of Minnesota St. Paul campus with 1.2 meters between each plant.

### Genotyping and Variant Detection

Tissue was collected from seedling leaves of the 74 Land Institute progeny, and adult leaf tissue was collected from the parents and all other progeny, with all tissue lyophilized prior to DNA extraction. Dual indexed genotyping-by-sequencing (GBS) libraries were created from genomic DNA using the restriction enzymes *SbfI* and *TaqI* and sequenced on 1.5 Illumina NovaSeq 6000 lanes (1×100 single-end reds). All extraction, library preparation, and sequencing occurred at the University of Minnesota Genomics Center.

Demultiplexed reads were mapped to *S. integrifolium* genomic contigs using BWA-MEM (Y. Brandvain, unpublished; Li, 2013). The “ref_map” pipeline of Version 2.5 of the “Stacks” variant calling software was then used for the detection of SNP loci (Catchen *et al.*, 2013). Loci that were missing in at least 30% of the population were excluded, as were loci missing in either parent. This resulted in 935 SNP markers.

### Linkage Map Construction

Linkage analysis and map construction were carried out in *JoinMap5* (Stam, 1993). First, 715 SNP markers showing significant (Chi-square test, P > 0.01) segregation distortion were removed. Markers were then grouped by LOD independence score, using a threshold score of five. Any groups that contained fewer than four markers were discarded. Map order and distances were then estimated using the maximum likelihood mapping algorithm, with default settings. One marker was removed from the end of linkage group three because it created a gap of more than 50 cM. Linkage group names were assigned based on estimated centimorgan length, with group one being the longest.

### Crossing design to determine self-incompatibility genotype

Of the 191 members of this mapping population planted in Minnesota, 84 were intermated for S-allele determination. To cross, capitula (compound flower heads) to be used as a female, or both as a male and female, were covered with a cotton bag prior to anthesis to prevent pollination. Capitula to be used exclusively as a male parent were covered with mesh bags at least one day prior to crossing to reduce contamination by insects depositing pollen from other plants. After stigma emergence, pollen from a male parent capitulum was collected into a container and dusted onto stigma with a pipe cleaner. Alternatively, on occasion a capitulum was removed and used to brush pollen directly onto a female parent. Each female capitulum was mated with only one male parent. The bag was then reclosed, harvested after senescence, and dried. For each capitula, the number of filled seeds and total number of seeds were then counted, with filled and unfilled seeds differentiated by visual and manual assessment. Seed set was then calculated as the ratio of filled seeds to total seeds.

Individuals were crossed in a structured design, which we have named the “connected small diallel design”, illustrated in Figure 1. This structure is based on groupings of four individuals that are mated in a diallel. These small diallels are then linked together through reciprocally mating single individuals. The purpose of this design was to maximize the amount of information that could be derived from the mating behavior of any given individual, while minimizing the number of matings for which it would need to be used. The design ensures that all individuals may be connected to one another through pairs of matings, allowing for the entire population to be used to predict the mating behavior each individual.

**Figure 1:**
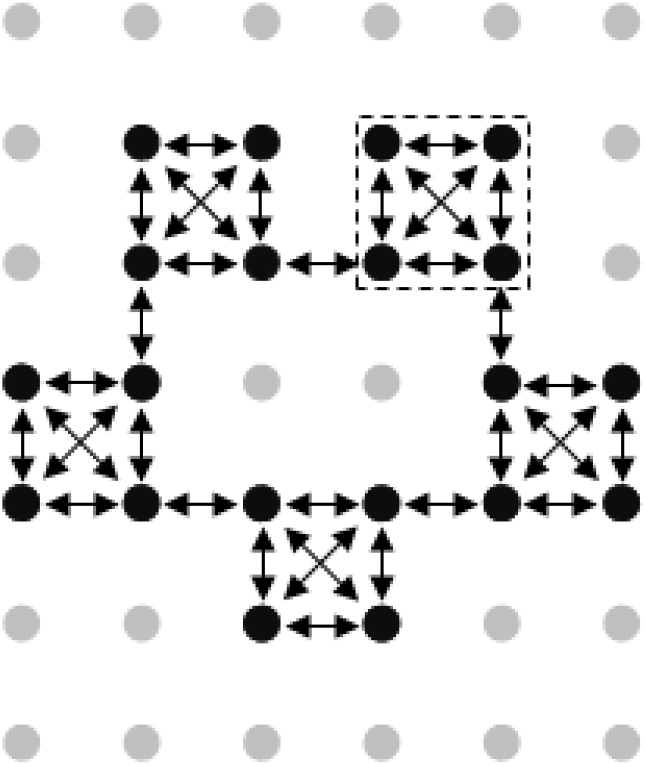
Visualization of the crossing design used in this experiment. Each circle represents one plant, with black circles representing plants that were used for crosses, and gray plants representing other members of the mapping population that were not selected. The dashed box represents one of the small diallels that formed the basis for the design—twenty of these were used for this experiment. Arrows represent matings, with each arrow pointing from the male parent to the female parent.

The design for this experiment included 20 small diallels. The design was implemented incompletely, with many of the recommended crosses not completed due to time or pollen availability constraints. To ensure that all individuals could be connected, more crosses connecting diallels were conducted than indicated in the design. Finally, several additional individuals were included in the study with only reciprocal crosses to one other individual. In total 268 crosses were performed between the 84 individuals, covering 138 different combinations of parents. Of these crosses, 126 combinations were crossed reciprocally, with both individuals used as male and female. One hundred and eighty-six crosses were conducted in 2018, and an additional 82 were conducted in 2019, primarily to increase the number of reciprocal crosses. In seven cases, multiple replications of a cross were performed. These observations were combined by summing the total number of filled and total seeds from each replication, and then calculating a seed set from the sums.

Three distinct but complementary methods were used to translate the results of the connected small diallel experiment into an S-locus map position. In all three methods, each mated pair of individuals is first determined to be compatible or incompatible, based on a threshold seed set value. A threshold value of 20% was selected for this experiment, based on previous observations of seed set in manually self-pollinated individuals (Reinert *et al*., 2020). Matings that resulted in a seed set value above 20% were considered compatible. These methods, along with their relative strengths and weaknesses, are described below. All methods were developed using the R language (R Core Team, 2019).

### Direct mapping with single-marker regression

The simplest method attempted to directly associate variation for compatibility with an SNP marker, without inferring the S-genotype of any particular individual. To accomplish this, a logistic regression was conducted following the formula S_i_ = M_ij_ + P_ij_ + M_ij_:P_ij_, where the success **S** of the ****i****th mating was predicted by the maternal genotype **M**, the paternal genotype **P**, and the interaction of those genotypes, all for the ****j****th marker. The calculated P-value of the maternal by paternal genotype interaction was then used to determine association with cross incompatibility, and thus the S-locus. Only biallelic markers were used in this approach, with homozygotes recoded as −1 or 1 and heterozygotes as 0. This approach only makes limited use of available genotype data, as each SNP marker is considered individually. Additionally, this approach could not be applied to species without genetic marker data. Finally, because it does not make inferences about the S-genotype carried by any particular individual or take potential dominance relationships into account, it is not able to fully leverage the advantages of the connected crossing design used in this study. Thus, this method’s usefulness is likely limited to confirming the results of other methods.

### Inference of self-incompatibility genotype for all individuals followed by QTL mapping of the inferred S-genotype

As an alternative approach to mapping the S-locus, we develop two methods to infer every individual’s S-locus genotype from its crossing behavior, then map this inferred phenotype with traditional QTL mapping software. Both methods assumed that the two parents of the population were each heterozygous at the S-locus and did not share any S-alleles with each other, resulting in a population with four distinct S-alleles and therefore four distinct S-genotypes. Each method is described in detail below. Code to replicate all methods may be found on GitHub (see data availability statement).

#### Approach 1: A hill-climbing algorithm

The first S-allele determination method used a simple hill-climbing algorithm to fit genotypes, given a user-generated set of dominance relationships among alleles. To start, the user hypothesizes a set of crossing relationships, dictating which of the sixteen possible pairings of genotypes will and will not be able to successfully mate. A random genotype is then assigned to each individual in the population. One arbitrarily selected individual is always set to a predetermined genotype to enable comparison of different runs of the algorithm. Each mated pair in the dataset is then scored using the user-supplied crossing relationships. A mismatch between the observed and predicted outcomes results in a score between 0.5 and 1, determined by the user. A match results in a score of one minus the previously mentioned score. Scores for the entire dataset are then summed. Next, the genotype of one individual is randomly changed, and the dataset rescored. If the total score decreases, then that set of genotypes is kept; if not, the algorithm returns to the previous position.

The cycle is repeated, stopping if 1,000 changes are attempted without decreasing the score. For this experiment, the algorithm was run 4,000 times for each set of dominance relationships, with the solution to the lowest scoring run considered optimal genotype assignments for a given dominance relationship. The lowest-scoring genotype assignments across all dominance relationships were then used for QTL mapping. We compared the results of 48 dominance relationships, each of which met three criteria: 1) individuals sharing both alleles must be incompatible, 2) individuals sharing no alleles must be compatible, and 3) there must be at least one combination of genotypes that was asymmetrical (compatible when one individual was used as a female and incompatible when the other was used as a female), as this was observed in the crossing data.

This method has both the advantage and disadvantage of being highly parameterizable. This makes it flexible, and gives the user a high degree of control, but may take many attempts to find the right combination of parameters to develop a solution. This method may be employed using a personal computer, however the reduction in parallelization necessary to achieve this may make it take too long to be useful. In addition, as a hill-climbing method there is no guarantee that this approach will find a global maximum.

#### Approach 2: A Markov Chain Monte Carlo algorithm

The second method employed sets of extreme gradient boosting decision trees with Markov chain Monte Carlo (MCMC) algorithms to infer S-locus genotypes. For each MCMC chain, an initial set of genotypes was created by first assigning a random heterozygous S-genotype to one random individual. The genotype of an individual that had been mated to the initial individual was then determined. If the cross was incompatible, the second individual’s genotype could share one or two alleles with the initial individual’s genotype, with equal probability. If the cross was compatible, the second individual’s genotype could share either zero or one allele with the initial plant’s genotype, with equal probability. The procedure was then repeated by choosing, at random, an individual that did not yet have a genotype assigned and that was crossed to the most recent individual assigned a genotype. If all individuals that were crossed to the most recently assigned individual had an assigned genotype, a random individual that did not yet have an assigned genotype was selected and the process started again until all individual were assigned genotypes.

Maternal and paternal S-locus genotypes were then treated as predictive variables with cross success used as a binary response variable. Constraints were placed on predictive variables so that matching maternal and paternal genotypes were not allowed to interact with one another, as crosses between individuals with matching genotypes should always be incompatible. At each step of an MCMC chain, model performance was measured as error of a logistic regression for classification using an extreme gradient boosting model with four-fold cross validation with 200 iterations and maximum tree depth of four splits, using the R package “xgboost” (Chen & Guestrin, 2016). A single genotype was altered at each MCMC chain step, using the same sets of probabilities used to construct the initial set. The newly proposed set of genotypes was accepted if the ratio of errors from the proposed genotype set to the former genotype set was greater than a randomly drawn number bound by zero and one.

Each MCMC chain, starting from a unique set of genotypes, was allowed to explore parameter space for 96 hours (~ 1 million steps). The set of genotypes that produced the smallest error from each chain was then used as the starting condition for hill-climbing algorithms. We used the same extreme gradient boosting model conditions described above, however at each step proposed genotypes were only accepted if they produced an error that was less than the previous set of genotypes. The hill-climbing algorithms each ran for 96 hours. For each chain, the set of genotypes that produced the smallest error, calculated as the percentage of crosses classified incorrectly, were recorded. A cross was predicted to be a success if the estimated logistic regression probability was greater than 0.5. The lowest error models were considered the best candidates for S-genotype determination. This method has the advantage of less dependence on user decisions, making it more repeatable and facilitating the exploration of a wider space of possible solutions. However, the resources required by this method mean that it generally cannot be employed on a personal computer.

#### Mapping the S-locus from inferred S-locus genotypes

Assigned alleles from the two inference methods were then used as phenotypes for QTL mapping, using the “qtl” R package (Broman *et al*., 2003), with the population treated as a four-way testcross. Marker genotypes were imputed using the “sim.geno” function, and QTL were identified using the “scanone” function to conduct interval mapping using the EM algorithm. Self-incompatibility allele was treated as a binary trait, with one allele from each parent arbitrarily coded as “1” and the other as “0”, and the two parental alleles mapped separately. For the MCMC method, all 96 of the inferred genotype sets were mapped, with the maximum LOD score produced by each set serving as a criterion to differentiate the sets. Eight genotype sets produced by the hill-climbing method were also mapped. Significance thresholds were determined independently for each set of genotypes used for mapping; only QTL with an error probability less than 5% were considered significant. This threshold ranged from a LOD score of 3.6 to 4.2. Finally, a Bayesian credible interval (similar to a confidence interval) around any identified QTL was calculated, using the “bayesint” function.

### Synteny with related species

To help elucidate the relationship between the *S. integrifolium* S-locus and other *Asteraceae*, the genomic contigs associated with the loci that comprise this linkage map were aligned to the annual sunflower (*Helianthus annuus*, variety ‘HA412’, version HOv1.1) (Badouin *et al*., 2017) and lettuce (*Lactuca sativa*, variety ‘Salinas’, version 7) (Reyes-Chin-Wo *et al*., 2017) genome assemblies using BLASTN (Altschul *et al*., 1990). Syntenic regions were then identified using the R package “syntR” (Ostevik *et al.*, 2020). For synteny analysis, all BLASTN alignments for a given *S. integrifolium* query sequence that had a bitscore greater than 90% of the maximum bitscore for that query sequence were used.

### Identification of candidate genes

For regions identified as syntenic to the *S. integrifolium* S-locus, gene annotations for sunflower were obtained from the genome assembly HA412v1 (Badouin *et al.*, 2017). InterPro taxonomy was searched for each gene in the region to identify plausible candidates by shared terms with S-genes in other species. The male and female determinants from *Brassica oleracea* (*Brassicaceae*) and *Petunia hybrida* (*Solanaceae*), as well as the female determinant of *Papaver rhoeas* (*Ranunculales*) chosen as representatives for comparison (The UniProt Consortium, 2019). In addition, defensin-like proteins were considered a proxy for the *Ipomoea trifida* (*Convolvulaceae*) male determinant (Rahman *et al.*, 2007). These species were selected because their self-recognition genes are relatively well characterized and they represent a broad range of Eudicot diversity, including both SSI and GSI systems. Any genes homologous to known CRPs were also considered potential candidates (Marshall *et al*., 2011). The best homologs for a subset of these candidates were then identified in the *S. integrifolium* reference transcriptome (Raduski *et al*., 2021), and their expression was measured in a previously-published set of whole-plant seedling transcriptomes from a diversity panel of 73 wild collected *S. integrifolium* accessions (Raduski *et al*., 2021). Expression was quantified in total transcripts per million (TPM), TPM as a percentile of total expressed genes, and TPM as a percentile of a reference set of conserved single copy genes, known as BUSCO (Simão *et al.*, 2015). We hypothesized that strong S-locus gene candidates would not be highly expressed in seedling vegetative tissue (Williams *et al*., 2014).

Additionally, the coding region sequence for the *B. oleracea* S-receptor kinase protein (Stein *et al*., 1991) was aligned to the lettuce and sunflower genomes using BLAST. These alignments were then compared to the lettuce and sunflower regions syntenic to the *S. integrifolium* S-locus.

## RESULTS

### Linkage Map

The linkage map contains 198 markers, spanning 1,049 centimorgans and divided into 7 linkage groups (Fig. 2, Table S1). This is consistent with the observation that *S. integrifolium* has seven chromosomes (Settle, 1967). The average distance between markers is 5.5 cM, with 29 gaps greater than 10 cM.

**Figure 2:**
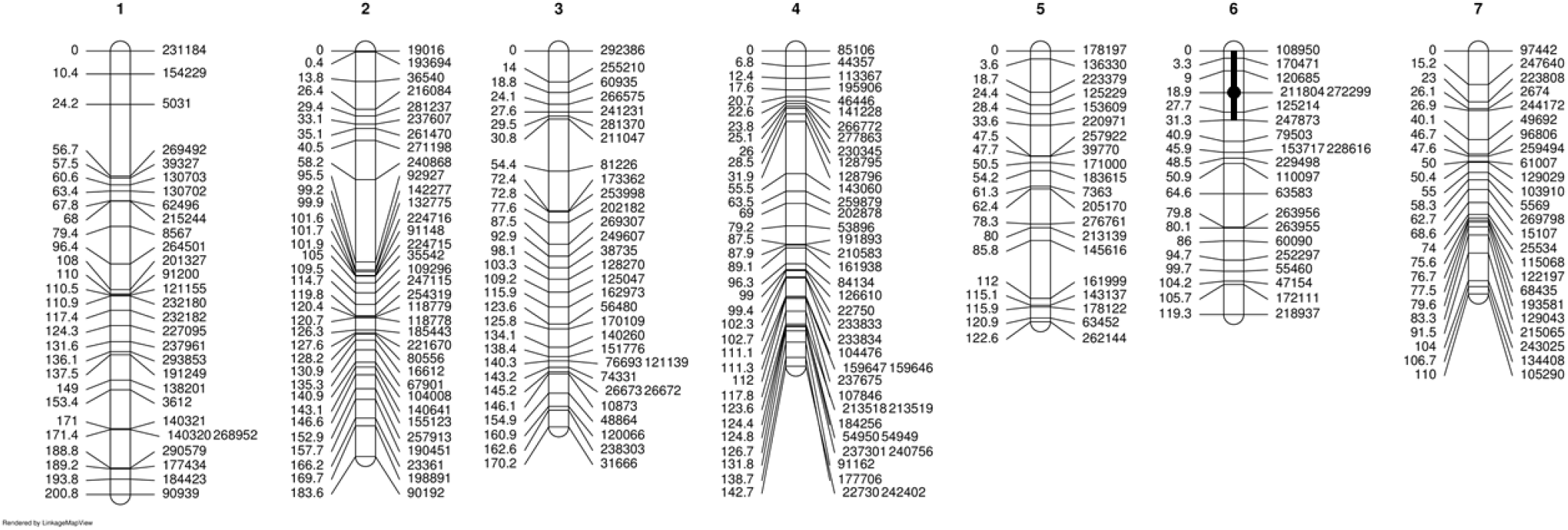
Linkage map for *Silphium integrifolium.* The black bar on linkage group 6 represents the putative S-locus QTL, with the circle showing the LOD peak and the extent of the bar showing a composite of the 95% Bayes credible intervals for several mapping methods. Map visualization was performed using “LinkageMapView” (Ouellette *et al*., 2018).

### S-locus mapping

Of the 268 crosses completed, 53% were incompatible, with a seed set value less than 20%, and 47% were compatible. These frequencies were not significantly different from an equal occurrence of compatible and incompatible crosses (df = 1, χ^2^ = 0.956, P = 0.328). This indicates there is a hierarchy of dominance between alleles in this population—if all alleles were codominant a ratio of 25% compatible to 75% incompatible would be expected. Of the 126 reciprocal pairs of genotypes crossed, 30% were compatible in both crossing directions (both individuals could be used as male and female), 34% were incompatible, and 36% were asymmetrical, or compatible when one individual was used as a female and incompatible when the other was used as a female. The presence of asymmetrical crosses indicates that the dominance relationship between some allele pairs differs from anthers to stigma. These observations confirm that *S. integrifolium* has a sporophytic SI system, as dominance relationships are not observed in gametophytic systems (Breton *et al*., 2014). Figure 3 shows the distribution of seed set values for reciprocal crosses, numerical data for all crosses may be found in Table S2.

**Figure 3:**
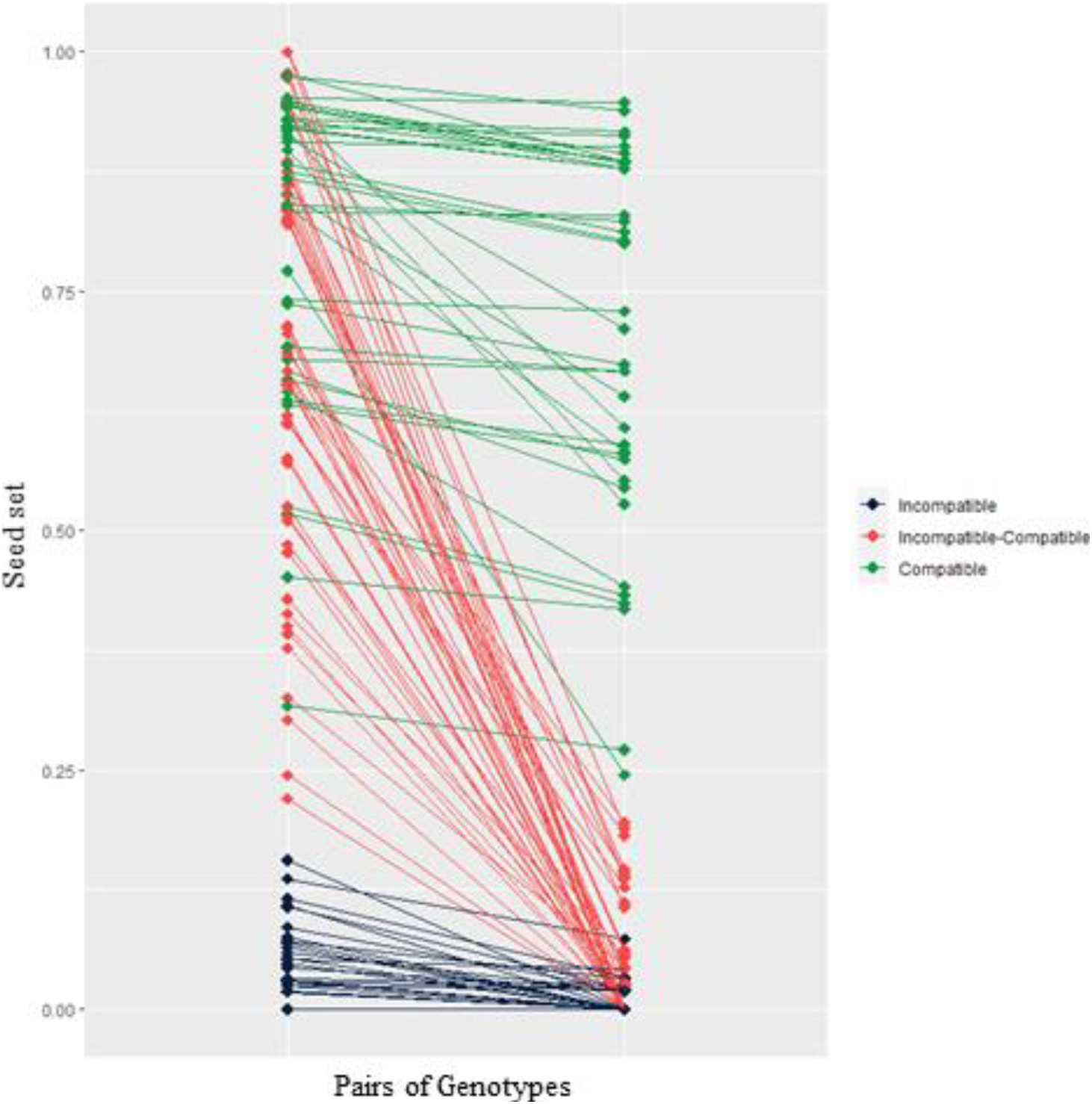
Distribution of seed set values for reciprocal matings (pairs of individuals mated, with each individual used as both a male and female). Each dot represents the seed set value of one plant, with lines connecting mated pairs. The individuals with the higher seed set value in the pair are placed on the left side of the chart.

#### Single marker regression for male/ female interaction

The interaction of male and female marker genotype was found to be a significant (P > 0.001) predictor of crossing success for four markers, located at two regions on the linkage map: markers “272299” and “120685” located at 18.9 and 9 cM, respectively, on linkage group six (P = 7.57 × 10^−8^, P = 1.46 × 10^−4^ df = 259 for both), and markers “185443” and “118778”, located at 126.33 cM and 120.7 cM, respectively, on linkage group two (P = 3.22 × 10^−5^, P = 3.31 × 10^−4^, df = 259 for both).

#### Hill-climbing algorithm method

Of the 48 dominance relationships tested using the hill-climbing algorithm, nine were used in QTL mapping. Of these, seven produced at least one significant LOD peak, all located on linkage group 6. The highest of these (LOD = 5.3) was located at 18.9 cM, and all but one of the other six were located between 9-31 cM.

#### MCMC method

Of the 192 genotype sets produced by the MCMC method (96 models, with a separate genotype set for each of the two parental alleles), 30 produced at least one significant LOD peak when used for QTL mapping, covering five linkage groups. Eleven of these genotype sets produce a significant LOD peak at either 18.9 or 27.7 cM on linkage group 6, and a further five were located elsewhere between 0-31 cM on linkage group 6. No other linkage group contained more than five of the 30 significant LOD peaks. The highest LOD peak produced by a single MCMC model was 7.56, located at 18.9 cM on linkage group six. This model also had the lowest error, and so we have selected it as the best result from the MCMC method. This set of genotype assignments is illustrated in Figure 4.

**Figure 4:**
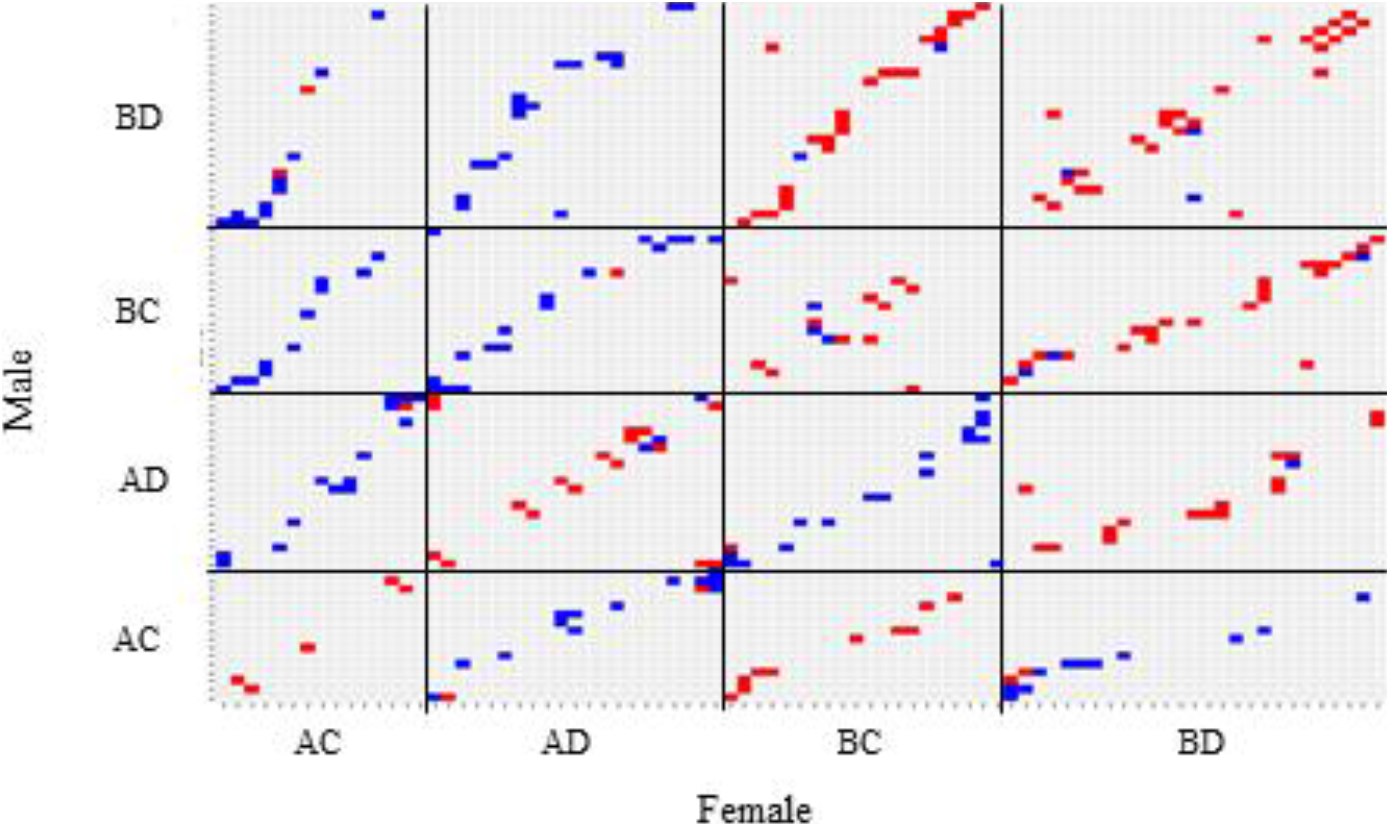
Representation of all possible matings that could have been completed for this study, with each square in the heatmap representing one pairing. Red or blue squares represent the crosses that were actually made, with red representing incompatible crosses, and blue representing compatible. Individuals are grouped by S-locus genotype, as assigned by the best MCMC genotype inference model. “A” and “B” refer to the S-alleles from one parent of the population, and “C” and “D” to the alleles from the other.

#### Consensus map location

Of the three methods used to map the S-locus, the MCMC method produced the strongest association between a genomic region and S-allele. The peak of this QTL was located at 18.9 cM on linkage group six, with a 95% probability Bayes credible interval from 0 to 27.7 cM. Broadly speaking, the results of the hill-climbing method agree with this region, although the credible interval for several of those results extend to 31 cM. Additionally, the most significant single-marker regression association between the interaction of male and female genotype with cross success was found at 18.9 cM. Taken together, we conclude that the *S. integrifolium* S-locus is located between 0 and 31 cM on linkage group six, with the marker closest to the locus likely located at 18.9 cM.

### S-locus synteny with related species

Synteny was found between this the putative *S. integrifolium* S-locus region and both the sunflower and lettuce genomes. In sunflower, synteny was found with two chromosomes—the *S. integrifolium* 0-18.9 cM region was found to align with chromosome 17, from 230.7 Mb to 262.67 Mb, and the 18.9-31cM region was found to align with chromosome 3, from 133.21 Mb to 149.9 Mb. In lettuce, the 0-9 cM region aligned with chromosome 4, from 18.13 Mb to 27.29 Mb, and the 18.9-31 cM region aligned with chromosome 9, from 144.19 Mb to 147.34 Mb. The syntenic region on sunflower chromosome 17 does not appear to overlap with the putative self-incompatibility breakdown QTL found on that chromosome by Gandhi *et al*. (2005), which we estimate was somewhere between 42 and 98 Mb, based on the alignment of SSR marker primer sequences to the sunflower genome assembly.

### Identification of candidate genes

Of the 642 genes located within the sunflower genomic regions syntenic with the *S. integrifolium* S-locus, 42 shared at least one InterPro term with either the male or female determinants of self-incompatibility in *B. oleracea* or *P*. *hybrida*, the female determinant of *P. rhoeas*, or the male determinant of *I. trifida*. Thirty of these genes were, like *B. oleracea* SRK, protein kinases, which is not a specific enough homology to infer a gene candidate. The other 12 genes were, like the female determinant in *P. hybrida*, F-box genes. In addition, one gene was similar to “STIG1”, a CRP. The expression of this gene, as well as two of the 12 F-box genes, were then measured in the 73 diversity panel seedling transcriptomes. One of these F-box genes was expressed in 65 of the seedling transcriptomes, with TPM as a percentile of BUSCO genes ranging from 5.4 to 79.5. The other was expressed in 42 of the seedlings, with TPM as a percentile of BUSCO genes ranging from 0.09 to 41.2. The STIG1-like gene was expressed in five of the seedling transcriptomes; expression varied widely but was always relatively low (Table 1).

**Table 1:** Expression of two S-gene candidates in a set of whole-plant seedling transcriptomes from a diversity panel of 73 wild collected *Silphium integrifolium* accessions. Expression is for *S. integrifolium* homologs of the named sunflower genes.

| Sunflower gene | Description | Plant ID | Transcripts per million (TPM) | TPM as percentile of all genes | TPM as a percentile of BUSCO genes |
| --- | --- | --- | --- | --- | --- |
| Ha3_00036110 | “Stig1-like” gene | C7 | 0.13 | 0.076 | 0.052 |
|  |  | D8 | 2.26 | 0.352 | 0.332 |
|  |  | E4 | 0.57 | 0.256 | 0.206 |
|  |  | E5 | 0.57 | 0.264 | 0.237 |
|  |  | H2 | 0.2 | 0.130 | 0.096 |
| Ha3_00036112 | serine/threonine protein kinase | B3 | 0.27 | 0.177 | 0.130 |
|  |  | B9 | 1.14 | 0.315 | 0.305 |
|  |  | E8 | 1.37 | 0.340 | 0.329 |
|  |  | H7 | 0.07 | 0.027 | 0.019 |

The best alignment for *B. oleracea* SRK in sunflower was found on chromosome 11, while the best alignment in lettuce was found on chromosome 7. Neither of these alignments are near any regions syntenic with the putative *S. integrifolium* S-locus in either species.

## DISCUSSION

### The described S-locus mapping methods are effective and efficient

This study represents one of the few successful identifications of a genomic region containing the S-locus in an *Asteraceae* species. This result is valuable for its own sake, as the comparison of syntenic genomic regions between *S. integrifolium* and other species may help to answer questions about the evolution of SSI systems in other *Asteraceae*. In addition, this study lays the groundwork for future efforts to clone the *S. integrifolium* S-locus.

In this experiment, we were able to predict the S-locus genotype for 84 individuals by conducting 268 crosses, with an average of 3.2 crosses per individual. This method did not require all 84 individuals to be crossed with a single tester, and so could be applied to a pre-existing full-sibling family without necessitating clonal propagation of any individuals. As a point of comparison, we estimate that the mapping of the S-locus in chicory, which combined a twenty-two plant diallel with testcrossing, required approximately 3,000 crosses (Gonthier *et al*., 2013). That approach, which also required the clonal propagation of tester genotypes, was used to assign an S-genotype to approximately 350 individuals (multiple replications of each cross were performed). The chicory mapping effort produced a more accurate determination of S-genotype and thus a narrower S-locus QTL region than our study, but with the trade-off of requiring more crosses per individual and the clonal replication of tester genotypes. Additionally, our method requires less time to complete, as testcrossing cannot be conducted until the diallel is completed, the results interpreted, and selected testers clonally propagated. Finally, our method does not require every cross to be replicated, as the analysis relies on inference methods that are relatively robust to error in the data.

Our method should be readily applicable to any other species that meet three criteria: 1) They possess an SSI system, 2) large full sibling families can be produced, and 3) individuals can readily be used as both a male and female parent in three to five matings. Among numerous other species, self-incompatible members of the genus *Helianthus* could be excellent candidates for this method and could help to confirm our results. Applying this method to phylogenetically diverse *Asteraceae* species could evaluate the conservation of the method of SSI in the family. Additionally, with some modifications, our framework could be used for GSI species.

### Evaluation of potential S-locus candidate genes

Of the 642 annotated or predicted genes present in the sunflower genomic regions syntenic to the putative S-locus, 43 show potential similarity to known S-genes. Of these, two groups of genes were considered particularly interesting, and are detailed below. It is important to note that because virtually nothing is known about the molecular mechanisms that underlie SSI in the *Asteraceae*, any discussion of potential candidate genes based on annotations or similarity to S-related genes in other systems is at best informed speculation. However, we see this as an important step towards identifying the genes that control SSI in the *Asteraceae*, and therefore believe it is worth pursuing.

One potential candidate is the sunflower gene Ha3_00036110, which is annotated as a putative member of the “Stigma Specific Protein 1” like, or STIG1-like, group of genes. In sunflower, this gene is found on the chromosome 3 region syntenic with the 18.9-31 cM segment of the putative *S. integrifolium* S-locus. In lettuce, the best BLAST alignment for Ha3_00036110 is on chromosome 9, within the region syntenic to the putative *S. integrifolium* S-locus, implying that this region may be conserved between the three species.

The best characterized member of this family, STIG1, encodes a small, cysteine-rich protein (CRP). In tomato (*Solanum lycopersicum*), this protein is primarily found in stigma exudate, and binds a pollen-specific kinase to promote pollen tube growth (Huang *et al*., 2014). In tomato, silencing either STIG1 or the pollen kinase resulted in slowed pollen tube growth and reduced seed set (Huang *et al*., 2014). More broadly speaking, as CRPs, STIG1-like genes are members of a gene class that also includes the male determinant of SSI in the *Brassicaceae* and the *Convolvulaceae*, and the female determinant of GSI in the genus *Papaver*. Based on this similarity to other SI genes, the function of STIG1, and the location of this gene within the sunflower genomic region syntenic to the *S. integrifolium* S-locus, we speculate that Ha3_00036110 is a signaling protein that controls the female determinant of SSI in *S. integrifolium*, with differential binding between different alleles of the STIG1-like protein and an unidentified pollen kinase resulting in compatibility or incompatibility. If this is correct, a possible candidate for this pollen kinase may be the gene Ha3_00036112, a serine/threonine protein kinase located 109 Kbp downstream from Ha3_00036110.

The best *S. integrifolium* homolog for Ha3_00036110 was only expressed in five of 73 seedling transcriptomes, and Ha3_00036112 was found to be expressed in four of 73 seedlings. The two genes were never expressed in the same individual, and expression was relatively low, never exceeding the 33rd percentile of BUSCO genes (Table 1). The low level of seedling expression for these genes further supports their candidacy, as S-gene expression is expected to be primarily limited to floral tissue (Williams *et al*., 2014). Expression in other tissues may still occur at low levels, either due to gene expression noise or as-yet uncharacterized uses for the gene product. For example, in tea (*Camellia sinensis*), an S-gene candidate was found to be expressed in leaf tissue at approximately 20% the level it was expressed in style tissue (Zhang *et al.*, 2016).

Another set of potential candidate genes are found on the sunflower chromosome 17 region syntenic with the 0-18.9 cM segment of the putative *S. integrifolium* S-locus. This region contains 12 F-box genes, resembling known gametophytic self-incompatibility (GSI) systems. For example, in *Petunia*, 17 tightly linked F-box genes serve as the male determinant of GSI (Williams *et al*., 2015). However, no genes in this sunflower region share any annotation terms with known S-RNAse genes (The UniProt Consortium, 2019), which serve as the female determinant in GSI systems. In addition, two of these genes that were arbitrarily selected as representatives were found to be expressed in more than half of available seedling transcriptomes, further diminishing the likelihood that they control the self-incompatibility response. Because so little is known about the molecular basis of the *Asteraceae* SSI response, these genes cannot be completely discounted, but they appear to be less likely candidates than the STIG1-like gene.

In addition to providing candidates for genes involved in *S. integrifolium* self-recognition, this study suggests that SRK-like genes are not involved in *S. integrifolium* SSI, as the best homologues for the *B. oleracea* SRK in the lettuce and sunflower genomes are not found in or near the regions of those genomes syntenic with the putative *S. integrifolium* S-locus. This finding adds evidence to the theory that specific genes that underlie *Asteraceae* SSI are different than those in the *Brassicaceae* (Allen & Hiscock, 2008).

### Implications for domestication and breeding

We expect the results of this study to facilitate the domestication of *S. integrifolium*. The availability of a genetic map, associated with particular restriction enzymes, supports the relatively inexpensive GBS genotyping of large numbers of progeny; both because these enzymes are now known to be useable in *S. integrifolium*, and because it is reasonably likely that some of the same loci would be recovered if these enzymes were applied to a different population. This may support the implementation of marker-based selection, marker-based pedigree development, and genome-wide selection. Additionally, this genetic map will assist in the anchoring and orientation of future genome assemblies.

Identifying a map location for the S-locus may also contribute to practical breeding efforts. If molecular marker data is routinely available for individuals within a breeding program, it may be possible to predict whether any two plants will be able to successfully mate, through the direct identification of particular S-alleles or ancestral haplotype blocks in the S-region. This information could save effort by excluding crosses that would not be successful. Perhaps more importantly, this would allow for the S-allele diversity of a given population to be monitored and maximized. It is likely that at least some *S. integrifolium* cultivars will take the form of synthetic populations, where a set of superior genotypes are intermated and their progeny form a distinct variety that may be reproduced for several generations. If a synthetic population is released that contains a low number of S-alleles, its long-term fecundity may be adversely affected by the limited number of individuals that are able to intermate. Alternatively, as *S. integrifolium* is known to express moderate to severe inbreeding depression for a number of traits (Price *et al.*, 2021), it is possible that S-allele characterization information could be used increase long-term productivity by limited mating among relatives for several generations within synthetic populations.

Overall, we anticipate that the availability of a genetic map and identification of the self-incompatibility locus will support efforts to domesticate *S. integrifolium* as a crop that will help to enable sustainable agricultural systems.

## ACKNOWLEDGMENTS

Funding for this work was provided by The Perennial Agriculture project in conjunction with The Land Institute and the Malone Family Land Preservation Fund, the United States Department of Agriculture’s National Institute of Food and Agriculture Grant no. 2019-67011-29607 to JHP, the Minnesota Department of Agriculture - Forever Green Agricultural Initiative, and NSF grant #1737827 Dimensions US-China to YB. The authors thank Shannon Lee Anderson, Karen Beaubein, and Jill Ekar for their help in completing the controlled crosses for this experiment. In addition, the authors thank Dr. Kevin Dorn for assistance in developing a GBS protocol, Dr. Adam Herman for bioinformatics advice, and Dr. Owen Beisel for inspiring the hill-climbing algorithm method. Finally, the authors acknowledge the Minnesota Supercomputing Institute (MSI) at the University of Minnesota for providing computing resources that contributed to the analysis of this study.

## AUTHOR CONTRIBUTIONS

The effort to develop a linkage map was initiated by KPS, DLVT, and YB, who also secured initial funding. DLVT created the mapping population, contributed text and visualization, and provided feedback in writing. JHP constructed the linkage map, initiated and designed the S-locus mapping work, and developed the HC method. ARR developed the MCMC method. JHP and ARR conducted data analysis and wrote the manuscript. KPS and YB provided supervision, and feedback on analysis, writing, and visualization.

## DATA AVAILABILITY

All code to replicate analyses may be found on GitHub, at https://github.com/UMN-BarleyOatSilphium/SilphiumSLocus. All sequence data may be accessed under BioProject PRJNA695552.

## SUPPORTING INFORMATION

Table S1: The linkage map in tabular form.

Table S2: All crossing data used in this experiment.

## References

Allen AM, Hiscock SJ. 2008. Evolution and phylogeny of self-incompatibility systems in angiosperms. In: Franklin-Tong VE, ed. Self-incompatibility in flowering plants: Evolution, Diversity and Mechanisms. Heidelberg, DE: Springer-Verlag, 73–101.

Allen AM, Thorogood CJ, Hegarty MJ, Lexer C, Hiscock SJ. 2011. Pollen-pistil interactions and self-incompatibility in the Asteraceae: new insights from studies of *Senecio squalidus* (Oxford ragwort). Annals of Botay 108(4):687–698.

Altschul SF, Gish W, Miller W, Myers EW, Lipman DJ. 1990. Basic local alignment search tool. J. Mol. Biol. 215:403–410

Badouin H, Gouzy J, Grassa CJ, Murat F, Staton SE, Cottret L, Lelandais-Brière C., Owens GL, Carrère S, Mayjonade B et al. 2017. The sunflower genome provides insights into oil metabolism, flowering and Asterid evolution. Nature 546(7656): 148–152.

Bod’ová K, Priklopil T, Field DL, Barton NH, Pickup M. 2018. Evolutionary pathways for the generation of new self-incompatibility haplotypes in a nonself-recognition system. Genetics 209(3):861–883.

Brennan AC, Harris SA, Hiscock SJ. 2006. The population genetics of sporophytic self-incompatibility in Senecio squalidus L. (Asteraceae): the number, frequency, and dominance interactions of S alleles across its British range. Evolution 60(2):213–224.

Breton CM, Farinelli D, Shafiq S, Heslop-Harrison JS, Sedgley M., Bervillé AJ. 2014. The self-incompatibility mating system of the olive (*Olea europaea* L.) functions with dominance between S-alleles. Tree Genetics & Genomes 10(4): 1055–1067.

Broman KW, Wu H, Sen Ś, Churchill GA. 2003. R/qtl: QTL mapping in experimental crosses. Bioinformatics 19:889–890

Camargo LEA, Savides L, Jung G, Nienhuis J, Osborn TC. 1997. Location of the self-incompatibility locus in an RFLP and RAPD map of *Brassica oleracea*. Journal of Heredity 88(1): 57–60.

Catchen J, Hohenlohe P, Bassham S, Amores A, Cresko W. 2013. Stacks: an analysis tool set for population genomics. Molecular Ecology 22(11):3124–40

Chen T, Guestrin C. 2016. XGBoost: A Scalable Tree Boosting System. In: Proceedings of the 22nd ACM SIGKDD International Conference on Knowledge Discovery and Data Mining (KDD’16). New York, NY, USA: Association for Computing Machinery, 785–794.

DeHaan LR, Van Tassel DL, Anderson JA, Asselin SR, Barnes R, Baute GJ, Cattani DJ, Culman SW, Dorn KM, Hulke BS et al. 2016. A pipeline strategy for grain crop domestication. Crop Science 56(3): 917–930.

Edh K, Widén B, Ceplitis A. 2009. Molecular population genetics of the SRK and SCR self-incompatibility genes in the wild plant species *Brassica cretica* (*Brassicaceae*). Genetics 181(3): 985–995.

Gonthier L, Blassiau C, Mörchen M, Cadalen T, Poiret M, Hendriks T, Quillet MC. 2013. High-density genetic maps for loci involved in nuclear male sterility (NMS1) and sporophytic self-incompatibility (S-locus) in chicory (*Cichorium intybus* L., *Asteraceae*). Theoretical and applied genetics 126(8): 2103–2121.

Harkness A, Brandvain Y. 2020. Nonself-recognition-based self-incompatibility can alternatively promote or prevent introgression. bioRxiv. doi: 10.1101/2020.09.29.318790

Harkness A, Goldberg EE, Brandvain Y. 2021. Diversification or collapse of self-incompatibility haplotypes as a rescue process. The American Naturalist. doi:10.1086/712424

Hiscock SJ. 2000. Genetic control of self-incompatibility in *Senecio squalidus* L.(*Asteraceae*): a successful colonizing species. Heredity 85(1): 10–19.

Hiscock SJ, Tabah DA. 2003. The different mechanisms of sporophytic self-incompatibility. Philosophical transactions of the Royal Society of London. Series B, Biological sciences 358(1434): 1037–1045.

Huang WJ, Liu HK, McCormick S, Tang WH. 2014. Tomato Pistil Factor STIG1 Promotes in Vivo Pollen Tube Growth by Binding to Phosphatidylinositol 3-Phosphate and the Extracellular Domain of the Pollen Receptor Kinase LePRK2. The Plant Cell 26(6): 2505–2523.

Fujii S, Takayama S. 2018. Multilayered dominance hierarchy in plant self-incompatibility. Plant reproduction 31(1): 15–19.

Gandhi SD, Heesacker AF, Freeman CA, Argyris J, Bradford K, Knapp SJ. 2005. The self-incompatibility locus (S) and quantitative trait loci for self-pollination and seed dormancy in sunflower. Theoretical and Applied Genetics 111(4): 619–629.

Larson S, DeHaan L, Poland J, Zhang X, Dorn K, Kantarski T, Anderson J, Schmutz J, Grimwood J, Jenkins J et al. 2019. Genome mapping of quantitative trait loci (QTL) controlling domestication traits of intermediate wheatgrass *(Thinopyrum intermedium*). Theoretical and Applied Genetics 132(8): 2325–2351.

Marshall E, Costa LM, Gutierrez-Marcos J. 2011. Cysteine-rich peptides (CRPs) mediate diverse aspects of cell–cell communication in plant reproduction and development. Journal of Experimental Botany 62(5): 1677–1686.

Kandel H, Hulke B, Ostlie M, Schatz B, Aberle E, Bjerke K, Eriksmoen E, Kraklau A, Effertz J, Rickertsen J. et. al. 2019. North Dakota Sunflower Variety Trial Results for 2019 and Selection Guide. Fargo, ND, USA: North Dakota State University Extension.

Koseva B, Crawford DJ, Brown KE, Mort ME, Kelly JK. 2017. The genetic breakdown of sporophytic self-incompatibility in *Tolpis coronopifolia* (*Asteraceae*). New Phytologist 216(4): 1256–1267.

Kubo K, Entani T, Takara A, Wang N, Fields AM, Hua Z. 2010. Collaborative non-self recognition system in S-RNase-based self-incompatibility. Science 330(6005): 796–799.

Li H. 2013. Aligning sequence reads, clone sequences and assembly contigs with BWA-MEM. arXiv:1303.3997 [q-bio.GN].

Price JH, Brandvain Y, Smith KP. 2021. Measurements of lethal and non-lethal inbreeding depression inform the *de novo* domestication of *Silphium integrifolium*. American Journal of Botany, in press.

Ostevik KL, Samuk K, Rieseberg LH. 2020. Ancestral Reconstruction of Karyotypes Reveals an Exceptional Rate of Nonrandom Chromosomal Evolution in Sunflower. Genetics 214(4): 1031–1045.

Ouellette LA, Reid RW, Blanchard SG, Brouwer CR. 2018. LinkageMapView—rendering high-resolution linkage and QTL maps. Bioinformatics 34(2): 306–307.

R Core Team. 2019. R: A language and environment for statistical computing. R Foundation for Statistical Computing, Vienna, Austria. URL https://www.R-project.org/.

Raduski AR, Herman A, Pogoda C, Dorn KM, Van Tassel DL, Kane N, Brandvain Y. 2021. Patterns of genetic variation in a prairie wildflower, *Silphium integrifolium*, suggest a non-prairie origin and untapped variation available for improved breeding. American Journal of Botany 108(1): 145–158.

Rahman MH, Uchiyama M, Kuno M, Hirashima N, Suwabe K, Tsuchiya T, Kagaya Y, Kobayashi I, Kakeda K, Kowyama Y. 2007. Expression of stigma-and anther-specific genes located in the S locus region of *Ipomoea trifida*. Sexual Plant Reproduction 20(2): 73–85.

Reinert S, Van Tassel D, Schlautman B, Kane N, Hulke B. 2019. Assessment of the biogeographical variation of seed size and seed oil traits in wild *Silphium integrifolium* Michx. genotypes. Plant Genetic Resources: Characterization and Utilization 17(5): 427–436.

Reinert S, Price JH, Smart BC, Pogoda CS, Kane NC, Van Tassel D, Hulke B. 2020. Mating compatibility and fertility studies in an herbaceous perennial Aster undergoing de novo domestication to enhance agroecosystems. Agronomy for Sustainable Development 40:27.

Reyes-Chin-Wo S, Wang Z, Yang X, Kozik A, Arikit S, Song C, Xia L, Froenicke L, Lavelle DO, Truco MJ et al. 2017. Genome assembly with in vitro proximity ligation data and whole-genome triplication in lettuce. Nature Communications 8(1): 1–11.

Rindos D. 1984. The Origins of Agriculture: An Evolutionary Perspective. Orlando, FL, USA: Academic Press.

Schiffner S, Jungers JM, Hulke BS, Van Tassel DL, Smith KP, Sheaffer CC. 2020. Silflower seed and biomass responses to plant density and nitrogen fertilization. Agrosystems, Geosciences & Environment 3:e20118.

Sedbrook JC, Phippen WB, Marks MD. 2014. New approaches to facilitate rapid domestication of a wild plant to an oilseed crop: example pennycress (*Thlaspi arvense* L.). Plant Science 227:122–132.

Settle WJ. 1967. The chromosome morphology in the genus *Silphium* (*Compositae*). The Ohio Journal of Science 67(1): 10–19.

Simão FA, Waterhouse RM, Ioannidis P, Kriventseva EV, Zdobnov EM. 2015. BUSCO: assessing genome assembly and annotation completeness with single-copy orthologs. Bioinformatics 31(19): 3210–3212.

Stam P. 1993. Construction of integrated genetic linkage maps by means of a new computer package: JoinMap. The Plant Journal 3: 739–744.

Stein JC, Howlett B, Boyes DC, Nasrallah ME, Nasrallah JB. 1991. Molecular cloning of a putative receptor protein kinase gene encoded at the self-incompatibility locus of *Brassica oleracea*. Proceedings of the National Academy of Sciences 88(19): 8816–8820.

Tomita RN, Fukami K, Takayama S, Kowyama, Y. 2004. Genetic mapping of AFLP/AMF-derived DNA markers in the vicinity of the self-incompatibility locus in *Ipomoea trifida*. Sexual Plant Reproduction 16(6): 265–272.

The UniProt Consortium. 2019. UniProt: a worldwide hub of protein knowledge. Nucleic Acids Research 47(D1): D506–D515

Van Tassel DL, DeHaan LR, Cox TS. 2010. Missing domesticated plant forms: Can artificial selection fill the gap? Evolutionary Applications 3(5–6): 434–452.

Van Tassel DL, Albrecht KA, Bever JD, Boe AA, Brandvain Y, Crews TE, Gansberger M, Gerstberger P, González-Paleo L, Hulke BS. et al. 2017. Accelerating Silphium Domestication: An Opportunity to Develop New Crop Ideotypes and Breeding Strategies Informed by Multiple Disciplines. Crop Science 57(3): 1274–1284.

Williams JS, Der JP, dePamphilis CW, Kao TH. 2014. Transcriptome analysis reveals the same 17 S-locus F-box genes in two haplotypes of the self-incompatibility locus of *Petunia inflata*. The Plant Cell, 26(7): 2873–2888.

Williams JS, Wu L, Li S, Sun P, Kao TH. 2015. Insight into S-RNase-based self-incompatibility in *Petunia*: recent findings and future directions. Frontiers in Plant Science 6:41.

Zhang CC, Wang LY, Wei K, Wu LY, Li HL, Zhang F, Cheng H, Ni DJ. 2016. Transcriptome analysis reveals self-incompatibility in the tea plant (*Camellia sinensis*) might be under gametophytic control. BMC Genomics 17:359.

